# Brain-to-brain synchrony during shared reminiscing predicts connection and memory in romantic couples

**DOI:** 10.64898/2026.09.11.751017

**Authors:** Jadyn S. Park, Lydia F. Emery, Yuan Chang Leong

**Affiliations:** Department of Psychology, University of Chicago, Chicago, IL 60637; Institute of Mind and Biology, University of Chicago, Chicago, IL 60637; Neuroscience Institute, University of Chicago, Chicago, IL 60637

**Keywords:** romantic relationships, social connection, memory, hyperscanning, fNIRS

## Abstract

Romantic partners often revisit relationship-defining memories together, reconstructing a shared account of their relationship through conversation. Such shared reminiscing can strengthen interpersonal connection by fostering alignment in thoughts, feelings, and perspectives, an experience known as shared reality. Here, we used functional near-infrared spectroscopy to record brain activity simultaneously from romantic couples during naturalistic, face-to-face conversations. Couples first discussed an unresolved conflict and then jointly recounted their relationship origin story, including how they first met and how their relationship developed. Shared reality decreased following the conflict discussion and returned to baseline levels after the origin story. While recounting their origin story, partners exhibited synchronized activity in the right medial frontopolar cortex, a region implicated in autobiographical memory and reasoning about close others. This synchrony was not observed at rest or during the conflict discussion. Greater medial frontopolar synchrony between partners during the origin story discussion was associated with higher post-conversation shared reality, closeness, and relationship satisfaction, even after accounting for baseline relationship quality. It also predicted better subsequent memory for the conversation, an association that was robust to controlling for the importance and familiarity of the memories being discussed. Together, these findings reveal how the interbrain alignment that emerges during shared reminiscing tracks both the connection partners experience and what they later remember. Our work bridges relationship science and social neuroscience through the naturalistic neuroimaging of live social interaction, and offers a window into the neural processes that accompany mutual understanding, social connection, and sharing memories in romantic relationships.

---

Romantic comedies often open with a “meet cute” – a charming first encounter between two characters that sets up a future romance. In real life, the first meeting is one of a series of relationship-defining memories that couples revisit and recount together over the course of their relationship. This shared reminiscing is not merely sentimental, but may also serve a relational function, helping partners maintain a clear and coherent sense of who they are as a couple (1). Indeed, couples who recount their shared past more positively, vividly, and with a stronger sense of “we” are more likely to stay together and report greater relationship satisfaction (2, 3).

When couples jointly recall relationship-defining memories, events are co-constructed through conversation, as partners selectively recall, elaborate, and negotiate what happened and what it meant (4–6). Through this process, couples may come to experience shared reality, the feeling that one’s thoughts, beliefs, and inner states are shared with another person (7, 8). In romantic relationships, perceptions of shared reality predict relationship satisfaction, closeness, and a sense of merged identity between partners (9–12). Joint reminiscing may be one pathway through which couples generate and renew this sense of shared reality, which then strengthens social connection and buffers against negative experiences (13).

In relationship science, shared reality has typically been measured by asking partners to report how much they experience shared thoughts, feelings, and perspectives about the world with their partner (14–16), or by having trained coders rate features of couples’ recorded conversations, such as how often they vocalize agreements or build on each other’s thoughts (10). Largely separate from this tradition, work in social and cognitive neuroscience suggests that alignment between minds may be visible at the neural level (17, 18). These studies have found that people who share interpretations of narratives exhibit temporally synchronous brain responses (19–22). Interbrain synchrony has also been shown to track social relationships, with friends having more similar brain responses when viewing the same content (23), and the degree of neural similarity between strangers while viewing videos predicts whether they will become friends (24).

Recent work has extended this idea to romantic couples, finding that partners exhibit greater neural similarity while watching relationship-relevant videos than random pairs (25). Hyperscanning paradigms, where neural activity is simultaneously recorded from both partners, have further shown that couples exhibit interbrain synchrony during cooperative tasks (26), handholding (27), and brief conversations (28, 29). To our knowledge, no study has examined interbrain synchrony during shared reminiscing. As such, it is unknown whether interbrain synchrony arises when couples jointly revisit their past, and whether such synchrony relates to the sense of shared reality and relationship quality that shared reminiscing is thought to foster.

Shared reality theory in relationships and the literature on interbrain synchrony have developed largely apart from one another. The current study brings these two literatures into conversation. We used functional near-infrared spectroscopy (fNIRS) to hyperscan romantic couples as they engaged in two face-to-face conversations (**Fig. 1**). Couples first discussed an unresolved conflict and then jointly recounted relationship-defining memories, enabling us to examine shared reminiscing following a relationship stressor. fNIRS tolerates the head movement inherent in live conversation and allows partners to speak, gesture, and respond to one another naturally (30, 31). Thus, our design situates the study of shared reality and interbrain synchrony in the naturalistic setting of free-flowing, face-to-face conversation.

**Figure 1.**
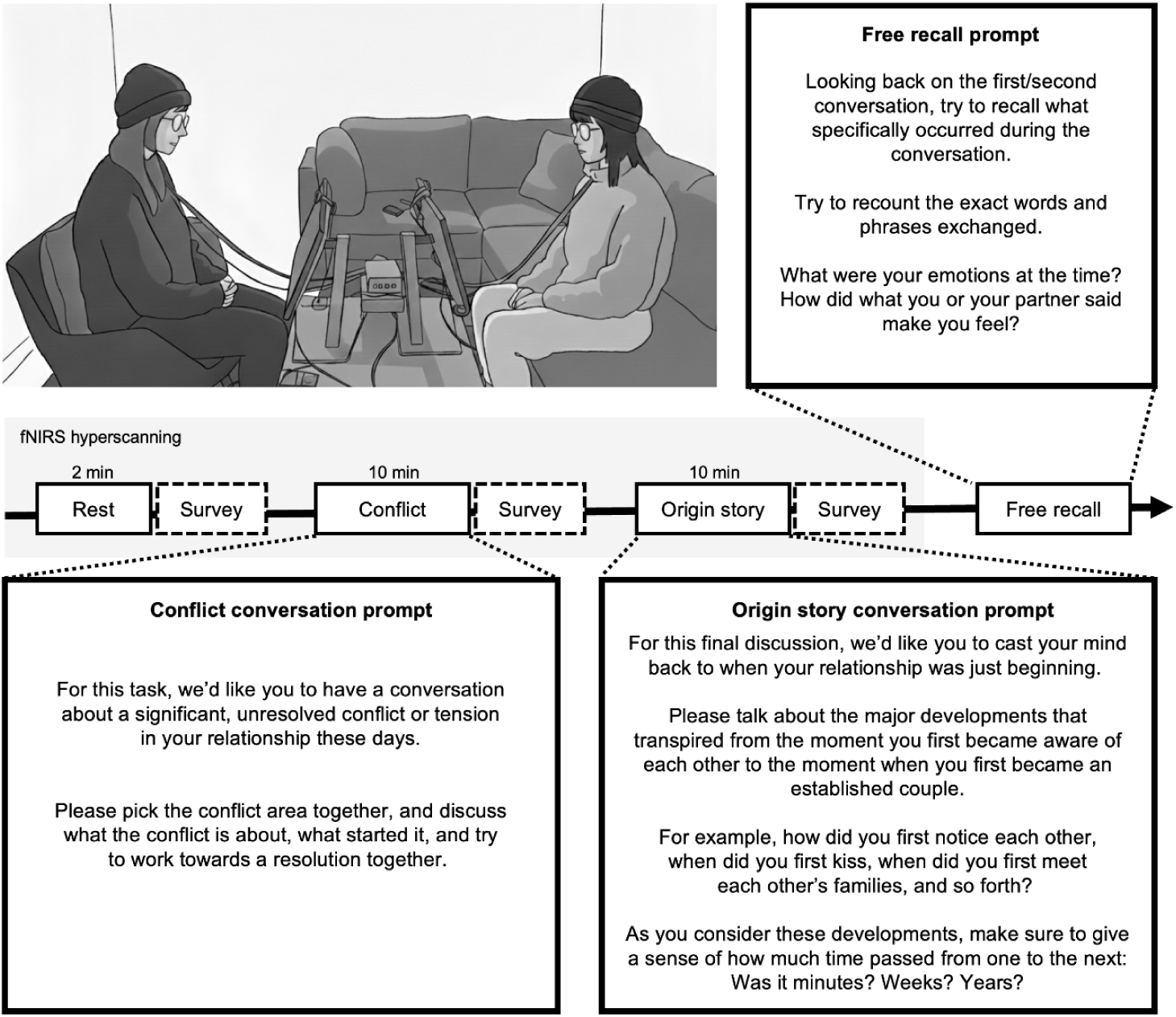
Schematic of experimental procedure. Couples sat across from each other while connected to a functional near-infrared spectroscopy (fNIRS) device. Participants completed two 10-minute conversations, each followed by a brief questionnaire. After both conversations, participants recalled each conversation from memory. Image (top left) created with Gemini from a staged photograph.

The first goal of our study was to test whether interbrain synchrony emerges during shared reminiscing and whether such synchrony predicts perceptions of shared reality and relationship quality following conflict. We focused on the prefrontal cortex, a region implicated in social cognition, memory retrieval, and the integration of self- and other-related information (32–36). Furthermore, recent studies in rodents have found that interbrain synchrony in the prefrontal cortex tracks social affiliative behavior (37–39). To quantify interbrain synchrony, we computed the inter-subject correlation (ISC) of neural time series between partners (40). We then tested whether ISC during shared reminiscing predicted post-conversation reports of shared reality, closeness, and relationship satisfaction, while controlling for baseline levels of these measures assessed prior to the conversations.

Our second goal was to examine whether interbrain synchrony during shared reminiscing predicted how well partners later remembered the conversation. Relationship-defining memories are revisited across the course of a relationship (41), and what partners retain from these retellings may become part of the narrative they carry forward (4). During joint retrieval, partners not only reactivate the memory of the original events, they also encode the conversation as a new episodic memory (42). Interbrain synchrony during shared reminiscing may index the extent to which partners are mutually engaged as the conversation unfolds (43–45). Such engagement may in turn support deeper encoding of the conversational episode for each partner, leading to more faithful subsequent memory for what was discussed. We therefore tested whether greater ISC during shared reminiscing predicted better individual recall of the conversation.

Together, these analyses offer a window into the neural dynamics that unfold between romantic partners as they jointly revisit the story of their relationship in a naturalistic conversation, and test whether the alignment of brain activity during these conversations is meaningful for social connection, what partners remember, and how they understand their relationship.

## Results

Forty-one romantic couples participated in our study (see **Fig. 2A** for demographic information and relationship status). All couples had been dating for at least 6 months, with a median relationship duration of 26 months (range = 6-293 months), and reported spending an average of 58 hours together per week (range=9-145 hrs/week; **Fig. 2B**). Prior to the experimental visit, participants completed the Generalized Shared Reality Scale (10), an 8-item measure assessing the extent to which they felt they shared thoughts, feelings, and beliefs with their partner. After each conversation, participants completed the same shared reality scale, rated how close and connected they felt to their partner during the discussion, and reported their current relationship satisfaction.

**Figure 2.**
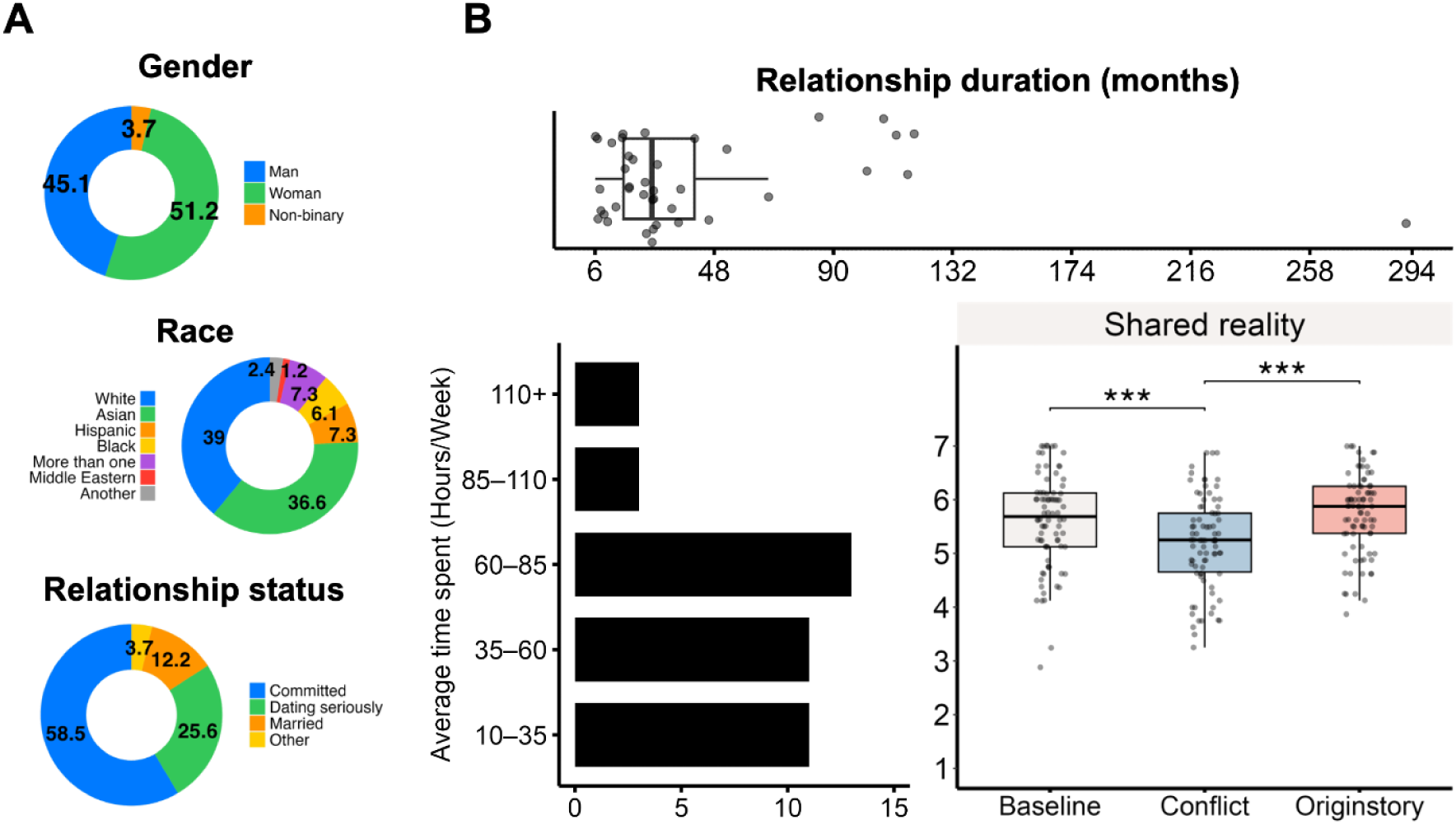
Sample characteristics and perceived shared reality across conversations. **(A)** Distribution of participants’ self-reported gender, race, and relationship status. Values indicate percentages. **(B)** Relationship duration (top), average time spent together per week (left), and shared reality at baseline and post-conversations (right). Individual data points represent the average value for each couple; boxplots indicate the median and interquartile range. Shared reality decreased following the conflict conversation and returned to baseline levels following the origin story conversation. ***p<.001.

Relative to baseline, participants reported a lower sense of shared reality following the conflict conversation (*baseline*: M=5.614, SD=.883; *conflict*: M=5.198, SD=.857; *t*(81)=-3.986, p<0.001), suggesting that discussing an unresolved conflict challenged participants’ sense of shared reality with their partner. Shared reality after the origin story conversation was higher than after the conflict conversation (*origin story*: M=5.73, SD=.735; *t*(81)=6.123, p<.001), and did not differ significantly from baseline (*t*(81)=1.325, p=.189; **Fig. 2B**). Participants also reported greater relationship satisfaction after the origin story conversation (M=6.609, SD=.715) than after the conflict conversation (M=6.451, SD=.918; *t*(81)=2.489, p<.05), and rated themselves as feeling closer and more connected to their partner during the origin story conversation (M=6.293, SD=.91) than during the conflict conversation (M=5.622, SD=1.263; *t*(81)=5.579, p<.001).

## Medial frontopolar cortex was synchronized between couples during shared reminiscing

We first asked whether romantic partners exhibited temporally aligned neural activity while jointly recounting relationship-defining memories. For each couple, we computed the channel-wise inter-subject correlations (ISC) between partners’ oxygenated hemoglobin (HbO) time series during the origin story conversation. We then calculated the median ISC across couples for each channel, and compared it against a null-distribution generated by recalculating the median ISC after randomly re-pairing each participant’s time course with that of a participant from a different couple (see **Methods**).

Romantic partners showed significantly greater ISC than randomly repaired participants in the right medial frontopolar cortex, an anterior midline portion of rostral prefrontal cortex (channel 13, median r=.08, p=.001, FDR *q*=.02; **Fig. 3A**; **Fig. 3B**). This region has been implicated in autobiographical memory and mentalizing about close or familiar others (32, 46, 47). Its recruitment during shared reminiscing is therefore consistent with the social-autobiographical nature of the task, in which partners jointly recall meaningful events from their relationship history. No fNIRS channels showed significant ISC during either the rest scan at the beginning of the experiment or during the conflict conversation (all *q*s > .05).

**Figure 3.**
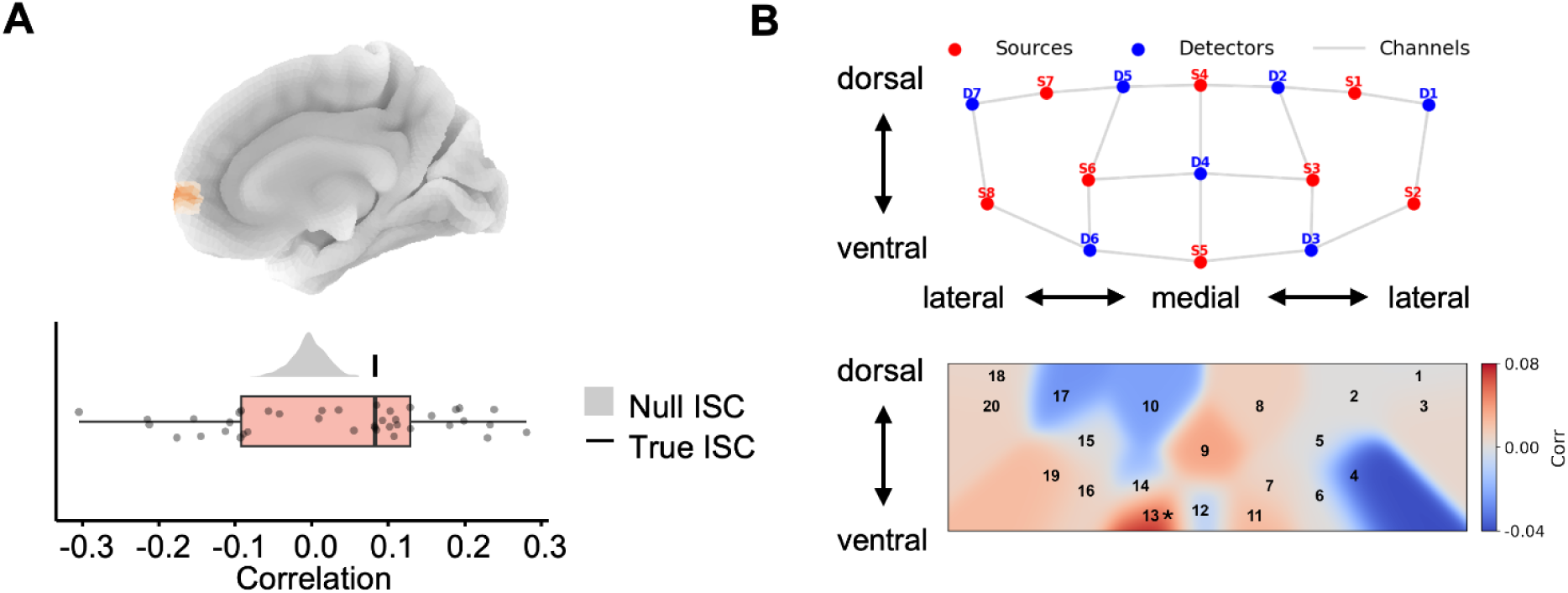
Neural time courses were synchronized in the right medial frontopolar cortex during the origin story conversation. **(A)** Cortical projection of the fNIRS channel with significant ISC (top). The observed median ISC, denoted as black vertical bar, is plotted against a null distribution generated by iteratively pairing each participant with a randomly selected individual who is not that participant’s partner. Points show ISC values for individual couples (bottom). **(B)** Layout of fNIRS optode placement, where red circles indicate sources, blue circles indicate detectors, and gray lines indicate channels (top). Heatmap shows ISC for each channel. Channel 13, marked with an asterisk, survived FDR correction at q=.02 (bottom).

## Medial frontopolar synchrony during shared reminiscing tracks shared reality, closeness, and relationship satisfaction

We then asked whether right medial frontopolar synchrony during the origin story conversation was associated with participants’ post-conversation self-reported measures. We ran these analyses using Bayesian mixed-effects regression models that accounted for the nesting of individuals within couples. All analyses controlled for relationship duration as well as baseline measures of relationship commitment and satisfaction collected during the intake survey (see **Methods**). Higher medial frontopolar ISC during the origin story conversation was associated with higher perceived shared reality during the conversation (*β*=.203, 95% HDI=[-.02, .41], p(*β*>0)=.965; **Fig. 4A**). The same results were observed when relationship duration, baseline commitment, and baseline satisfaction were excluded from the model (*β*=.217, 95% HDI=[-.02, .48], p(*β*>0)=.958).

**Figure 4.**
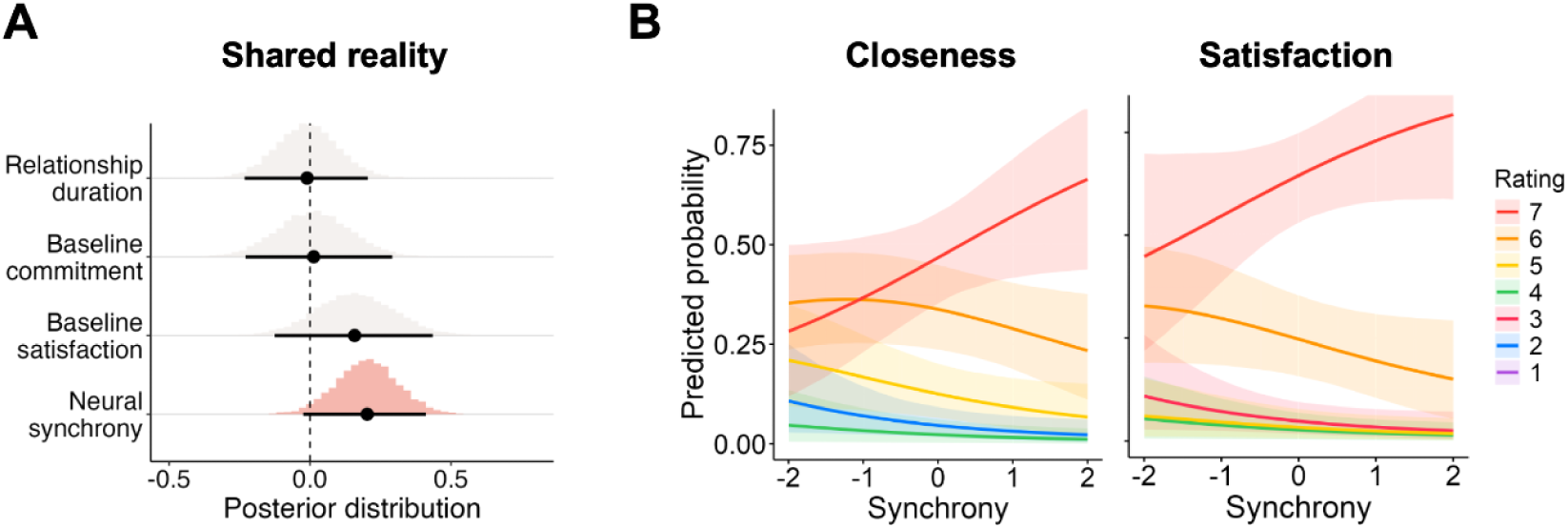
Medial frontopolar synchrony during shared reminiscing tracks shared reality, closeness, and relationship satisfaction. **(A)** Posterior distribution of the regression coefficient when predicting shared reality from medial frontopolar synchrony, estimated using a Bayesian mixed effects model. The 95% highest density interval (HDI) is indicated by the bold horizontal line, with the median indicated by the black circle. (**B**) Predicted probabilities from Bayesian ordinal logistic mixed effects models predicting closeness and relationship satisfaction from medial frontopolar synchrony. The x-axis shows interbrain synchrony in standardized units; the y-axis shows the model-predicted probability of each response category on the 7-point scale. Each colored line corresponds to one rating category (1–7), and shaded bands indicate 95% HDI.

We then tested whether medial frontopolar ISC was associated with participants’ ratings of closeness and relationship satisfaction. As these outcomes were each assessed with single-item rating scales, we fit Bayesian ordinal logistic mixed-effects regression models to account for the ordered categorical nature of the responses (48). Higher medial frontopolar ISC was associated with greater closeness during the origin story conversation (*β*=.43, median OR=1.53, 95% HDI=[.91, 2.26], p(OR>1)=.974) and higher post-conversation relationship satisfaction (*β*=.41, median OR=1.51, 95% HDI=[.86, 2.34], p(OR>1)=.952; **Fig. 4B**). The same pattern of results was observed when the analyses were repeated with Bayesian linear mixed-effects regressions (**Figure S1**), and when the ordinal models were rerun without relationship duration, baseline commitment, and baseline satisfaction as covariates (see **Figure S2**).

Across four separate models, relationship duration and baseline measures of relationship quality were themselves not associated with medial frontopolar synchrony (*Relationship duration*: *β*=-.151, 95% HDI=[-.47, .19], p(*β*<0)=.815; *Shared reality: β*=.02, 95% HDI=[-.23, .29], p(*β*>0)=.556; *Satisfaction: β*=.073, 95% HDI=[-.21, .35], p(*β*>0)=.697*; Commitment: β*=-.133, 95% HDI=[-.39, .12], p(*β*<0)=.852; **Figure S3**). These results suggest that our findings were not due to couples with higher relationship quality exhibiting higher medial frontopolar synchrony during the origin story discussion. Instead, medial frontopolar synchrony appears to reflect interaction-specific processes during shared reminiscing that tracked with how connected partners felt afterwards.

## Medial frontopolar synchrony during shared reminiscing predicts subsequent memory of the conversation

Next, we asked whether medial frontopolar synchrony during shared reminiscing predicted participants’ subsequent memory. We first examined whether synchrony varied with the degree to which the origin story was viewed as central to who they were as a couple, a feature that could plausibly shape partners’ engagement during the conversation and their later recall. Couples who rated the origin story as more central exhibited higher medial frontopolar ISC during the conversation (β=.60, 95% HDI=[.17, 1.06], p(β>0)=.997, OR=1.82), suggesting that interbrain synchrony was strongest when partners jointly recounted memories that were important to their shared identity.

To examine the relationship between medial frontopolar synchrony and memory of the conversation, we used Google’s Universal Sentence Encoder (49) to convert the conversation transcripts and participants’ subsequent recall of the conversation into embedding vectors that capture semantic similarity between texts. For each participant, we quantified recall fidelity as the cosine similarity between the vector encoding the conversation transcript and that encoding the participant’s recall (**Fig. 5A**). This approach allowed us to quantify the semantic overlap between the recalled and the original conversational content, providing a continuous and context-sensitive measure of memory (50–52).

**Figure 5.**
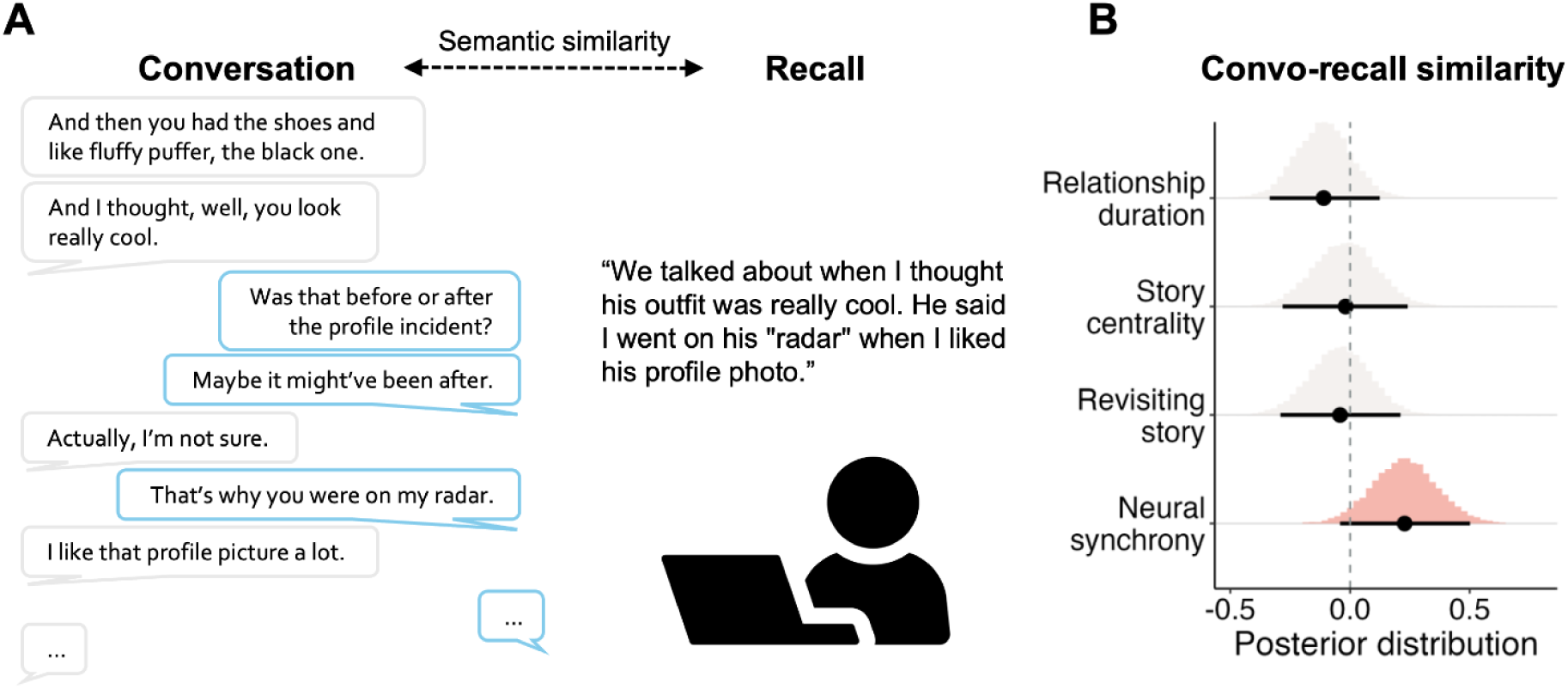
Medial frontopolar synchrony during origin story conversation predicts subsequent memory for the conversation. **(A)** Schematic depicting the origin story conversation and subsequent memory recall from an example couple. Each participant’s recall and the transcript of the origin story conversation were converted into text embeddings, and recall fidelity was calculated as the cosine similarity between the two embedding vectors. **(B)** Posterior distribution of the regression coefficient when predicting shared reality from medial frontopolar synchrony, estimated using a Bayesian multilevel model. The 95% HDI is indicated by the bold horizontal line, with the median indicated by the black circle. Higher medial frontopolar synchrony during the origin story conversation was associated with higher subsequent recall fidelity of the conversation.

Participants with higher medial frontopolar synchrony during the origin story conversation recalled the conversation with higher fidelity (β=.24, 95% HDI=[-.02,.48], p(β>0)=.966). This association was robust to controlling for the perceived centrality of the origin story to relationship identity, how frequently participants reported revisiting the story, and the duration of the relationship (*β*=.23, 95% HDI=[-.05, .49], p(*β*>0)=.953; **Fig. 5B**). Together with the findings above, these results suggest that medial frontopolar synchrony not only captures perceived shared reality and connection, but may also index the extent to which participants encode the experience.

## Exploratory analyses examining conversational dynamics and medial frontopolar cortex synchrony

We next explored whether observable conversational dynamics were related to medial frontopolar synchrony during shared reminiscing. We were particularly interested in turn-taking behaviors, as they are a fundamental feature of dyadic interaction and have been linked to self-disclosure and interpersonal rapport (53). We therefore coded two relevant turn-taking behaviors: backchannels, brief vocal acknowledgement of engagement and attention (e.g., “mmhmm”, “yeah”), and interruptions, defined as taking over the conversational floor. We coded these behaviors during both the conflict and origin story conversations, allowing us to ask whether turn-taking dynamics differed across conversational contexts and whether dynamics during either conversation was associated with neural synchrony during shared reminiscing.

Couples used more backchannels (*conflict*: 17.8%, SD=11%; *origin story*: M=8%, SD=6%; t(36)=6.549, p<.001) and interruptions (*conflict*: M=11%, SD=6%; *origin story*: M=5%, SD=3%; t(36)=5.176, p<.001) during the conflict discussion relative to the origin story discussion. Medial frontopolar cortex synchrony was not related to the number of backchannels during the conflict (β=-.07, 95% HDI [-.37, .22], p(β<0)=.696) or origin story discussions (β=-.05, 95% HDI [-.32, .22], p(β<0)=.638). Similarly, medial frontopolar synchrony was not associated with interruptions during the origin story discussion (β=.05, 95% HDI=[-.20, .32], p(β>0)=.666). In contrast, the number of interruptions during the conflict conversation was negatively associated with medial frontopolar synchrony during the origin story discussion (β=-.37, 95% HDI [-.62, -.11], p(β<0)=.996). In other words, couples who interrupted each other more when discussing an unresolved conflict were subsequently less likely to become neurally aligned when jointly recalling their relationship origin story. This association was robust to controlling for baseline measures of relationship commitment and satisfaction (*β*=-.43, 95% HDI=[-.67, -.17], p(*β*<0)=.998), as well as post-conflict shared reality (β=-.42, 95% HDI=[-.67, -.16], p(β<0)=.998).

## Discussion

Romantic couples often return to the memories that define who they are together. They revisit them on anniversaries, retell them to friends, and reconstruct them through acts of joint retrieval. In the present study, we used fNIRS hyperscanning to examine the neural dynamics that unfold as romantic couples engaged in shared reminiscing during a live, face-to-face conversation. We found that couples exhibited interbrain synchrony in the right medial frontopolar cortex while jointly recounting relationship-defining memories. Couples who showed greater medial frontopolar synchrony during shared reminiscing reported greater shared reality, closeness, and relationship satisfaction and later recalled the conversation with greater fidelity. Together, these findings suggest that shared reminiscing is accompanied by neural alignment between romantic partners, and that this alignment is meaningful for both the connection partners experience in the moment and what they later remember from the interaction.

Our findings extend the study of shared reality in close relationships by showing that the sense of sharing thoughts, feelings, and perspectives with a partner is reflected not only in what people report and how they behave, but also in the temporal alignment of neural activity. Prior studies have often relied on self-report measures (14–16) or behavioral coding of recorded interactions (10). These approaches have been central to establishing that shared reality matters for close relationships (9, 15), but are less well suited to measure alignment that unfolds during a social interaction. By measuring neural activity simultaneously from both partners during natural conversation, the present study captures shared reality as an interpersonal process that emerges over time. We do not suggest that interbrain synchrony is itself shared reality. Rather, the measure may track with psychological processes that support shared reality (17, 18, 54, 55). Consistent with this interpretation, interbrain synchrony has been previously shown to be associated with joint attention (56, 57), affective alignment (58–61), and arriving at a shared interpretation (62, 63).

During the origin story conversation, partners were not simply retrieving parallel versions of past events. They needed to access episodic memories, model their partner’s perspective on those events, and negotiate a coherent shared account (64, 65). These demands—retrieving autobiographical memory, mentalizing, and integrating narratives—have each been associated with the medial frontopolar cortex (32, 46, 47, 66–68). Medial frontopolar synchrony may emerge when partners become cognitively and affectively in step while they listen, recall, elaborate, and revise the story together. That this synchrony emerged during the origin story conversation but not during rest or the preceding conflict discussion suggests that medial frontopolar synchrony was not simply a general feature of being together or speaking with a partner. Instead, medial frontopolar synchrony may be more likely to emerge when partners coordinate autobiographical retrieval and perspective taking around a cooperative effort to construct a shared account. The right lateralization of our results is also consistent with prior evidence that emotional and self-relevant autobiographical retrieval recruits right rostral medial prefrontal regions (66, 69, 70), though evidence for hemispheric specialization in autobiographical memory has been inconsistent (71, 72), and this aspect of the finding should be interpreted cautiously.

Greater medial frontopolar synchrony during the origin story conversation was associated with greater perceived shared reality, closeness, and relationship satisfaction. These findings connect the literature on interbrain synchrony with accounts of shared reality in close relationships, which propose that relationships are strengthened when partners experience their thoughts, feelings, and perspectives as aligned (12, 14). One alternative explanation is that this association reflects pre-existing relationship quality, such that happier couples tend to exhibit higher interbrain synchrony. This account appears less likely, however, because baseline shared reality, satisfaction, and commitment did not predict medial frontopolar synchrony, and the association between medial frontopolar synchrony and post-conversation outcomes remained credible after adjusting for these baseline relationship quality measures. Thus, medial frontopolar synchrony appears to capture an emergent feature of the conversation, rather than reflecting stable differences in relationship quality.

Our findings should not be taken to imply that greater interbrain alignment always corresponds to stronger social connection. Indeed, a recent fMRI hyperscanning study using a *Fast Friends* paradigm found that friends became less neurally aligned over the course of the conversation, whereas strangers became more aligned (73). Among strangers, greater divergence was associated with more positive conversation outcomes. These findings suggest that the relationship between interbrain alignment and social connection may depend on relationship context and conversational goals. In the *Fast Friends* paradigm, where conversation partners get to know one another through structured prompts, neural divergence may reflect exploration and novelty that makes the conversation more engaging and rewarding. In our task, partners jointly reconstruct a shared autobiographical narrative, for which alignment may reflect the coordination of memories and perspectives. Taken together, these findings raise the possibility that effective social coordination involves knowing when to align and when to diverge.

Shared reminiscing not only reactivates memories of past events, but also creates a new episodic memory of the conversation that can later be recalled and integrated into partners’ shared understanding of their relationship. Medial frontopolar synchrony was also associated with how well partners remembered the conversation. This association remained credible after accounting for the perceived centrality of the origin story and how frequently partners had revisited it, suggesting that interbrain synchrony was not merely driven by the familiarity or personal importance of the material. Rather, higher synchrony may reflect periods of mutual engagement and coordinated attention that support deeper encoding (74–78). This interpretation builds on prior fMRI studies showing that intersubject correlation among individuals viewing the same movie scene predicts better subsequent memory for that scene (51, 79). Whereas that work examined separate participants exposed to the same stimulus, the present findings extend this association to live interaction, in which partners jointly shape the content they later remember.

In exploratory analyses, we asked whether observable turn-taking dynamics were associated with medial frontopolar synchrony. Couples who interrupted one another more during the preceding conflict discussion showed lower medial frontopolar synchrony during the origin story conversation. One possibility is that a disruptive conflict discussion carried over into the next interaction (80, 81), making it harder for partners to establish shared reality during the origin story discussion. Another possibility is that interruptions reflect a more general communication style, such that couples who more often talk over one another are also less likely to align with each other during the origin story conversation. The present study was not designed to distinguish between these explanations, particularly because the conflict discussion always preceded the origin story conversation. Future studies that counterbalance conversation order or assess turn-taking across multiple interactions could determine whether interruptions disrupt subsequent neural alignment or index a broader communication pattern within the couple.

One important limitation is that our study is correlational and cannot establish whether interbrain synchrony contributes causally to shared reality, connection, or memory. Experimental approaches that manipulate neural alignment across interacting partners could eventually address this question (82). In mice, synchronizing optogenetic stimulation of the medial prefrontal cortex across animals increased affiliative social interaction, providing evidence that experimentally induced interbrain synchrony can influence social behavior (38, 39). Recent advances in noninvasive multibrain stimulation methods provide the opportunity to test related hypotheses in humans by applying temporally coordinated stimulation to both members of a dyad and examining its effects on neural synchrony and psychological outcomes (83–85). Real-time hyperscanning neurofeedback may offer another complementary approach by training dyads to increase or decrease their neural alignment during social interaction (86–88). Our findings identify the medial frontopolar cortex as a promising target for causal manipulations. Future studies can take advantage of causal approaches to test whether increasing or decreasing interbrain synchrony fosters or disrupts social connection and memory for a jointly experienced interaction.

Altogether, our study provides evidence from humans engaged in naturalistic, face-to-face interaction that interbrain synchrony emerges as romantic partners jointly recall their shared past. These findings suggest that interbrain synchrony may reflect the processes through which partners co-construct mutual understanding and weave together a shared narrative of their relationship. More broadly, jointly revisiting the past may help couples sustain a sense of “us,” offering insight into the neural and interpersonal processes through which shared memories strengthen romantic bonds.

## Materials and Methods

### Participants

Forty-one romantic couples (82 individuals; age=20-55 years; 51.2% women, 45.1% men, 3.7% non-binary) were recruited from the greater Chicago community. To be eligible, participants had to have been dating their partner for at least 6 months. Seven couples reported that they were in a same-gender relationship. Thirty-nine percent of the sample identified themselves as White, 36% as Asian, 6% as Black, and 7% as more than one race. More than half of the participants reported that they were in a committed long-term partnership (58.5%), a quarter reported they were dating seriously (25.6%), and 12% reported they were married. Experimental procedures were approved by the University of Chicago Institutional Review Board, and all participants provided informed consent prior to the study. Participants were compensated with $40 (per person) for their time. One couple was excluded due to fNIRS device failure, resulting in a final sample of 40 couples (80 individuals) for analysis.

## Intake questionnaire

Prior to the experimental visit, participants completed an online intake survey that included demographic information as well as the following measures. All measures used 7-point Likert scales, unless otherwise noted.

### Generalized Shared Reality

Shared reality was measured using an 8-item scale that measures the extent to which individuals believe they and their partner share the same thoughts and beliefs about the world (10). At intake, participants completed the cross-situational version of the scale (e.g., “We typically share the same thoughts and feelings about things”). During the experiment, participants completed the interaction-specific version (e.g., “During our interaction, we shared the same thoughts and feelings about things”) (α=.85, M=5.61, SD=.88).

### Commitment

Commitment (e.g., “I want our relationship to last for a very long time”) was measured using the 7-item commitment subscale of the Investment Model Scale (89) (α=.83, M=6.61, SD=.55) .

### Satisfaction

Satisfaction (e.g., “I feel satisfied with our relationship”) was measured using the 5-item satisfaction subscale of the Investment Model Scale (89) (α=.78, M=6.39, SD=.58).

## Experimental design

After both members of each couple completed the intake survey, they were invited to the lab for a 90-minute experimental session. Participants sat facing each other across a table in a dimly lit room, with monitors placed in front of them for instructions (**Fig. 1**). The experiment consisted of three “runs” while participants underwent fNIRS recording. Each run was followed by a brief questionnaire.

In the first run, participants completed a 2-minute resting-state task in which they were instructed to close their eyes and relax. The second run consisted of a 10-minute conflict conversation in which participants discussed a recent, unresolved conflict in their relationship. The third run consisted of a 10-minute “origin story” conversation in which participants recalled how they first met and reflected on their relationship trajectory. After each conversation, participants completed the interaction-specific Generalized Shared Reality scale (10) (e.g., “During our interaction, we thought of things at the exact same time.”), single-item measures of closeness (“I felt close and connected to my partner during this discussion”) and satisfaction (“Right now, I feel satisfied with our relationship”). After the origin story conversation only, participants provided an additional single-item measure of story centrality (“This story is central to who we are as a couple”).

Following both conversations, participants were separated into different rooms and asked to recall each conversation from memory by typing their responses on a laptop.

## fNIRS recording and analysis

### fNIRS data acquisition

During the conversations, participants were scanned using fNIRS. The fNIRS data were collected using a NIRSport2 fNIRS device (NIRx Medical Technologies) with a sampling rate of 10.1725 Hz at wavelengths of 760 and 850 nm. The fNIRS device layout consisted of 20 channels composed of 8 source optodes and 7 detector optodes using the unambiguously illustrated (UI) 10/10 external position system (90). Raw fNIRS data were preprocessed in MATLAB using custom scripts based on the Homer2 package (91). Detailed preprocessing procedures are described in the **Supplementary Text**, which included correction for hemodynamic lag, exclusion of noisy channels, bandpass filtering, and motion correction. The oxygenated hemodynamic time courses were z-scored separately for each run and each participant.

### Inter-subject correlation analysis

For each channel, we calculated the inter-subject correlation (ISC) for each dyad by computing the Pearson correlation between partners’ HbO time courses, where a higher ISC indicates greater neural synchrony. To assess statistical significance, we used a permutation approach. In each permutation, every participant was randomly paired with another participant who was not their actual partner, and the correlation was computed for each randomly paired dyad. The median correlation across all randomly paired dyads were then calculated. This procedure was repeated 1,000 times to create a null distribution. We then calculated the p-value by counting the number of permutations where the “null” correlation was more extreme than the observed correlation. This approach minimized assumptions about the underlying signal distributions while preserving the temporal structure of the data.

## Conversation transcripts and memory performance

Audio recordings of the conversations were transcribed using WhisperX (https://github.com/m-bain/whisperx), specifically the large-v2 model, which was installed on a local server to ensure participant privacy and confidentiality. The transcripts were then manually cleaned to correct for typos, speaker labels, utterance timing, and any other discrepancies with the original conversations.

Utterances from the cleaned transcripts and post-conversation memory recall responses were then converted to sentence embeddings using Google’s Universal Sentence Encoder (49). Four couples were excluded from the memory analysis due to language translation (2 Chinese speaking couples) and audio quality issues (2 couples). To calculate the semantic similarity between the memory recall and the overall conversation, embeddings from the conversation utterances were first averaged, representing the overall semantic space of the conversation, and cosine similarity with the recall embeddings was then computed. Higher cosine similarity values reflected greater semantic alignment between the recall and the conversation.

## Bayesian multilevel models

To assess the relationship between interbrain synchrony and relationship-related outcomes, we used Bayesian mixed effects models with individuals nested within couples and random intercepts defined at the couple-level. Effects were estimated using linear models for multi-item composite outcomes and ordinal logistic models for single-item ordinal outcomes (see **Supplementary Text** for detailed model specifications). All models were implemented using the *brms* package (version 2.21.0) in *R* version 4.3.2, and were specified with the random seed “123” to ensure reproducibility. Weakly informative priors were implemented to support model convergence while still following the posterior distributions to be predominantly shaped by the data. Each model was run with four Markov Chain Monte Carlo (MCMC) chains, each with 4,000 samples, of which the first 1,000 samples were discarded as burn-in. The remaining samples from all chains were concatenated to form the posterior distribution of each parameter. Posterior distributions were summarized by reporting the estimated posterior medians and 95% highest density intervals (HDIs). For linear models, we considered the effects to be credible if more than 95% of the posterior distribution was either greater than or less than zero. For ordinal logistic models, posterior draws were exponentiated to obtain odds ratios (OR), and we considered effects to be credible if more than 95% of the posterior of the OR was greater than 1.

## Author Contributions

J.S.P., L.F.E., Y.C.L. designed research;

J.S.P. performed research;

J.S.P. analyzed data;

J.S.P., L.F.E., Y.C.L. wrote the paper.

