## Supplemental Materials for "Brain-to-brain synchrony during shared reminiscing predicts connection and memory in romantic couples"

This PDF file includes:
Supplementary Figures 1-3
Supplementary Text
Supplementary References

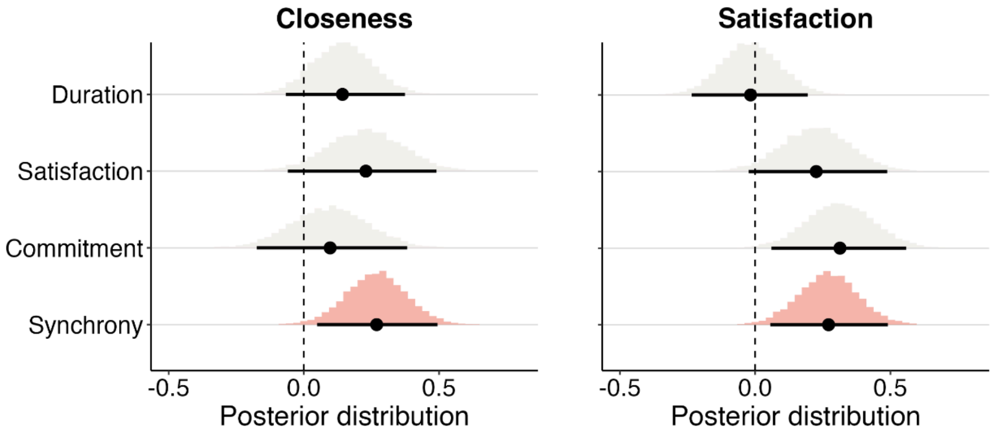


**Supplementary Figure 1. Medial frontopolar synchrony is associated with post-conversation closeness and satisfaction.** Posterior distribution of the regression coefficient when predicting closeness and satisfaction from medial frontopolar synchrony, estimated using a Bayesian linear mixed-effects model. The 95% highest density interval (HDI) is indicated by the bold horizontal line, with the median indicated by the black circle. These models included relationship duration, baseline levels of satisfaction and commitment as covariates.

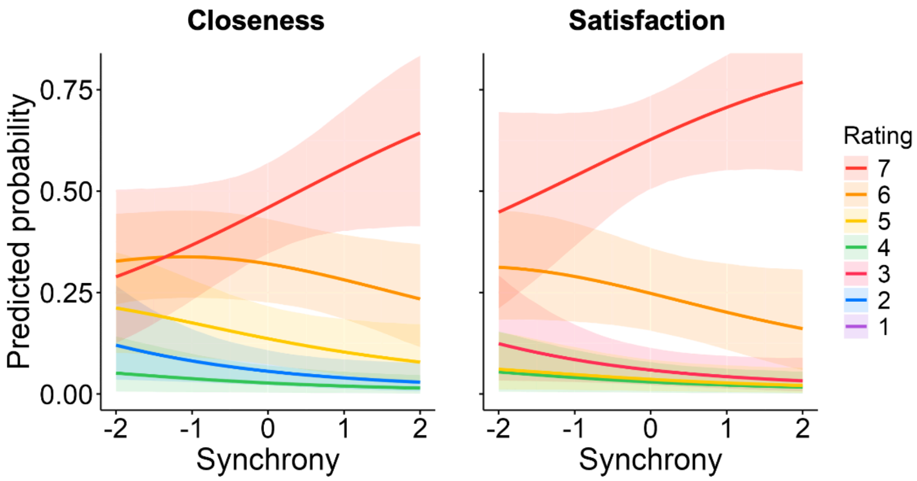


**Supplementary Figure 2. Medial frontopolar synchrony is associated with post-conversation closeness and satisfaction.** Predicted probabilities from Bayesian ordinal logistic mixed effects models predicting closeness and relationship satisfaction from medial frontopolar synchrony. These models did not include relationship duration, baseline satisfaction, and baseline commitment as covariates. The x-axis shows interbrain synchrony in standardized units; the y-axis shows the model-predicted probability of each response category on the 7-point scale. Each colored line corresponds to one rating category (1-7), and shaded bands indicate 95% HDI.

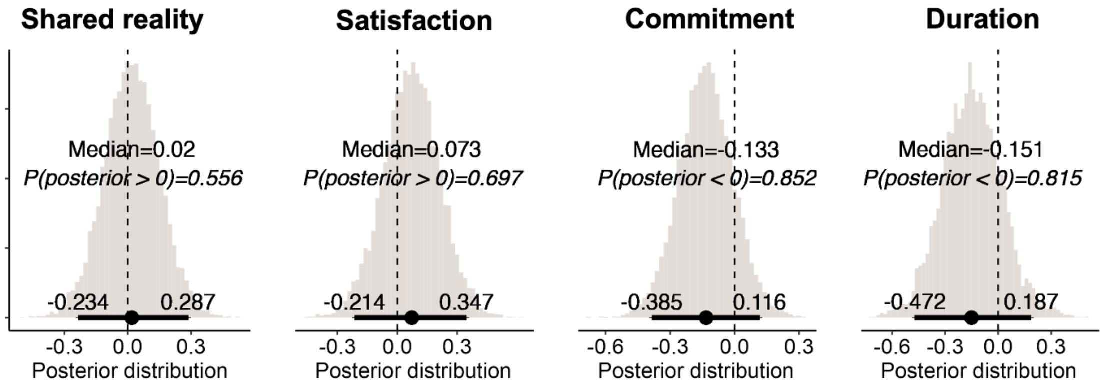


**Supplementary Figure 3. Baseline levels of relationship quality are not associated with medial frontopolar synchrony.** Posterior distribution of the regression coefficient for the association between baseline relationship quality and medial frontopolar synchrony, estimated using separate Bayesian linear mixed-effects models for each predictor. The 95% highest density interval (HDI) is indicated by the bold horizontal line, with the median indicated by the black circle.

### Supplementary Text

#### fNIRS data preprocessing and visualization

***Preprocessing.*** Raw fNIRS data were preprocessed in MATLAB using custom scripts based on the Homer2 package (Huppert et al., 2009). First, raw light intensity data were aligned across participants based on task triggers, with a 4.5 second shift applied to account for hemodynamic lag. Then, channels were classified as unusable if detector saturation occurred for longer than 2 seconds or if the light intensity signal resembled white noise. Specifically, the spectral characteristics of the signal were quantified using quartile coefficient of dispersion (QCoD), where channels with a QCoD less than 0.2948 were classified as unusable. The light intensity data were then converted to optical density, and bandpass filtered from 0.005 to 0.15Hz to remove low frequency drift and potential physiological signals (Lyu et al., 2024). Motion artifacts were identified based on changes in optical density exceeding five standard deviations or an absolute change in signal amplitude of two within a one-second interval. Principal components analysis-based motion correction was then applied with a variance threshold of 0.80. Optical density values were then converted to changes in oxygenated (HbO), deoxygenated (HbR) and total (HbT) hemoglobin concentrations using the Modified Beer Lambert Law. Finally, the HbO time series were z-scored across time.

***Spatial localization.*** The 3D coordinates of each channel were obtained in their native space. The MNI coordinates of five anatomical fiducials (Nasion, Nz; Inion, Iz; Central, Cz; Left auricular, AL; Right auricular, AR) were used as landmarks to derive transformation matrix from native space to MNI space. This transformation was then applied to the native-space coordinates of the remaining channels to obtain their corresponding MNI coordinates.

#### Bayesian multilevel models

***Linear models.*** For each subject $i$ and couple $j$, the observed outcome $y_{i,j}$ is assumed to follow a normal distribution, where:

$$y_{i,j} \sim Normal(\mu_{i,j}, \sigma)$$

$\mu_{i,j}$ denotes the expected outcome for subject $i$ and couple $j$, and $\sigma$ denotes the residual standard deviation of the outcome.

$\mu_{i,j}$ is modeled as a function of fixed intercept and fixed slope for the predictors with couple-specific random intercepts:

$$\mu_{i,j} = \beta_0 + \alpha_{couple[j]} + \beta_1 x_{i,j}$$
$$\beta_0 \sim Normal(0, 1)$$
$$\beta_1 \sim Normal(0, 1)$$
$$\alpha_{couple[j]} \sim Normal(0, \sigma_{couple})$$
$$\sigma_{couple} \sim Exponential(1)$$
$$\sigma \sim Exponential(1)$$

***Ordinal logistic regression.*** For each subject *i* and couple *j*, the observed outcome $R_{i,j}$ is assumed to follow a categorical distribution, where:

$$R_{i,j} \sim Categorical(\boldsymbol{p}_{i,j})$$
$$\boldsymbol{p}_{i,j} = \{p_{i,j1}, p_{i,j2}, \dots p_{i,jK}\}$$

The vector $\boldsymbol{p}_{i,j}$ denotes the probabilities of the response falling into each of the $K$ response categories (e.g., for a 7-point Likert scale, $K$=7).

The cumulative probabilities were modeled using logit link function,

$$logit(P[R_{i,j} \leq k]) = \kappa_k - \phi_{i,j}, k = 1, \dots, K - 1$$

where $k$ indexes the $K - 1$ thresholds ($k = 1, \dots, K - 1$), $\kappa_k$ denotes the $k$th threshold, and $\phi_{i,j}$ denotes the linear predictor:

$$\phi_{i,j} = \alpha_{couple[j]} + \beta_1 x_{i,j}$$
$$\kappa_k \sim Normal(0, 1)$$
$$\beta_1 \sim Normal(0, 1)$$
$$\alpha_{couple[j]} \sim Normal(0, \sigma_{couple})$$
$$\sigma_{couple} \sim Exponential(1)$$

The priors for these models were selected to be weakly informative. For fixed effects, we used Normal(0, 1) priors as we had no strong prior belief about the size or the direction of the effect. For standard deviation parameters, we used Exponential(1) priors to constrain values to be positive and assign greater probabilities for smaller standard deviations without strongly constraining the model (McElreath, 2016).

### Conversational prompts

Before each conversation, the prompts were read aloud to the participants by the experimenter. During the conversation, the prompts remained present on the screen in front of them. After 9 minutes, participants were presented with a notice stating, “You have one minute left.”

***Conflict conversation.*** For this task, we’d like you to have a conversation about a significant, unresolved conflict or gension in your relationship these days. Please pick the conflict area together, and discuss what the conflict is about, what started it, and try to work towards a resolution together.

***Origin story conversation.*** For this final discussion, we’d like you to cast your mind back to when your relationship was just beginning. Please talk about the major developments that transpired from the moment you first became aware of each other to the moment you first became an established couple.

For example, how did you first notice each other, when did you first kiss, when did you first meet each others’ families, and so forth? As you consider these developments, make sure to give a sense of how much time passed from one to the next: Was it minutes? Weeks? Years?

### Behavioral coding of interruptions and backchannels

Backchannels and interruptions were manually coded by trained research assistants who watched the video recordings of the conversations. For each utterance in the cleaned conversational transcript, research assistants indicated whether the utterance was followed by a backchannel and/or an interruption. Backchannels were defined as verbal signals of engagement and understanding (e.g., “mmhmm”, “yeah”), whereas interruptions were defined as instances in which their partner took over the conversational floor. These turn-taking signals were coded as binary variables for each utterance (1 = present, 0 = absent), irrespective of the number of instances following that utterance. For example, multiple consecutive backchannels (“yeah, yeah, yeah”) were treated as a single occurrence. Then, for each participant, the proportions of backchannels and interruptions were calculated separately as the number of utterances followed by each type of turn-taking signal, divided by the total number of utterances spoken by that participant in a conversation.
